# A cortical semantic space integrating fractions and integers

**DOI:** 10.64898/2026.03.24.713850

**Authors:** Daniela Valério, Samuel Debray, Alireza Karami, Maxime Cauté, Nicolás Gravel, Stanislas Dehaene

## Abstract

How does the human brain represent the meaning of abstract symbols? Some theories postulate the existence of semantic spaces where concepts occupy positions that reflect their conceptual relationships. In the number domain, psychological evidence suggests that integers are represented along a mental number line which, with education, integrates higher-level number concepts such as fractions. To test this hypothesis, we recorded whole-brain 7T fMRI responses to integer and fraction symbols during a magnitude comparison task. Consistent with predictions, we found both behavioral and neural numerical distance effects. Activation vectors in intraparietal, inferior temporal, prefrontal, hippocampal, and parahippocampal cortices formed a two-dimensional semantic space organized by numerical magnitude and domain (fractions versus integers). Gaussian fits revealed a topographic map of numerical preferences in the anterior inferior parietal cortex, common to both domains. Our results suggest that, in educated adults, a joint semantic map integrates fractions and integers and supports symbolic magnitude representation and comparison.

## Introduction

“*God made the integers, all else is the work of Man*” - Leopold Kronecker.

The issue of how the human brain represents the meaning of complex and abstract concepts is a challenging problem. A plausible hypothesis is that concepts are mapped onto internal conceptual spaces or “semantic maps” (Gardenfors, 2000). Within the high-dimensional space provided by the activity of millions of neurons in relevant areas of the brain, each concept would be represented as a specific point or vector, with semantic relationships encoded by the proximity between those vectors, constituting a neural implementation of the concept of second-order isomorphism (Shepard & Chipman, 1970). This view is supported by the discovery that, in both brain-imaging and multiple single-cell recordings, the major dimensions of semantic variations of words induce gradients of neuronal activation on the surface of the cortex (Deniz et al., 2019; Franch et al., 2025; Huth et al., 2016). However, while major semantic axes have been found to distinguish broad domains of knowledge, such as animate versus inanimate entities, whether a similar map-like organization exists for abstract concepts *within* a given domain, at the level of individual words, has received fewer attention (though see e.g. Borghesani et al., 2019; Mason et al., 2021; Mason & Just, 2016; Viganò & Piazza, 2020).

Here, we tackle this issue by studying a specific domain of semantic competence, central to human education: the cerebral representation of numerical symbols, including intuitive ones – such as the integers 0-5 – as well as more abstract forms which are acquired through education – fractions such as 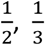, or 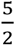. Understanding how these numbers relate to one another and what they mean are fundamental components of human cognition that lie at the foundation of scientific reasoning. The perception of concrete numerosities – that is, the number of items in a set - and their ratios appears early in life (Izard et al., 2009; McCrink & Wynn, 2007) and is consistently observed across individuals, regardless of formal education (McCrink et al., 2013; Pica et al., 2004). Even the magnitude zero, understood as the quantity of a set with no elements, can be grasped by monkeys and humans (Barnett & Fleming, 2024; Kutter et al., 2024; Nieder, 2016). However, through education and the acquisition of symbolic systems, humans developed the capacity to extend the concept of number far beyond those intuitions of small integers. Most seven-year-olds struggle with the notion that there might be numbers between 1 and 2, while, for college students, this idea seems obvious, because education expanded their concept of number to include fractions, decimals, and real numbers. The expansion of the number field, including the invention of negative and complex numbers, represents a uniquely human invention that provides new mathematical tools for modelling the world. Yet, how and where such abstract symbolic mathematical concepts are represented in the brain, and what kind of semantic space they form, remains poorly understood.

For integers, it is well established that individuals in Western cultures tend to associate smaller numbers with the left side of space and larger numbers with the right, the SNARC effect (Spatial-Numerical Association of Response Codes; Dehaene et al., 1993). Moreover, two numbers are more easily discriminated as the distance between them increases – a phenomenon known as the distance effect (Dehaene et al., 1990; Moyer & Landauer, 1967). Together, these findings support the idea that integers are mentally represented along a one-dimensional semantic map, a spatial continuum known as the mental number line (Dehaene et al., 1993; Dehaene & Changeux, 1993). Constructing this representation requires extracting the quantity associated with a number independently of its format – whether it appears as a word (“five”), a number (“5”), or a set of items (“five dots”). This process is typically attributed to the Approximate Number System (ANS), which is thought to involve the bilateral intraparietal sulci (IPS) as a primary cortical site (Barnett & Fleming, 2024; Dehaene et al., 2003; Piazza et al., 2007; Pinel et al., 2001; Walsh, 2003), although other potential candidates have also been proposed, including inferior temporal cortex and hippocampus (Cai et al., 2023; Kutter et al., 2023, 2024; Nelli et al., 2023; Pinheiro-Chagas et al., 2018). Interestingly, a similar representation principle may underlie fraction processing, as numerical magnitude (i.e., how large a fraction is) also plays a central role in understanding fractions but see also Bonato et al., 2007). Consistent with this idea, a distance effect has been observed during fraction processing in both behavioral (Kallai & Tzelgov, 2009; Kalra et al., 2020; Meert et al., 2010; Schneider & Siegler, 2010) and brain-imaging studies (Ischebeck et al., 2009; Jacob & Nieder, 2009; Mock et al., 2018), and a SNARC effect has also been found with fractions (Toomarian & Hubbard, 2018).

Here, our goal is to understand how symbolic fractions are represented both neurally and behaviorally and whether this representation aligns with a mental number line. We hypothesize that, in participants with sufficient education, a number line based on magnitude might integrate the two domains of integers and fractions. This is because, with education and expertise, individuals learn various strategies that help them place numbers correctly along a magnitude continuum (Cauté & Dehaene, 2025). If this is the case, we should expect to observe, at the behavioral level, a distance effect and an internal representational geometry (revealed through multidimensional scaling, MDS) organized according to numerical magnitude, independently of the number’s original domain (integers or fractions). This would be a first source of evidence for a shared number line between fractions and integers. At the neural level, using representational similarity analysis (RSA; Kriegeskorte et al., 2008), we expect to observe graded BOLD responses such that numbers closer on the mental number line show more similar neural responses. Additionally, if neurons putatively tuned to similar magnitudes cluster together on the surface of the cortex, we expect to find cortical regions where voxels tuned to symbolic numbers are topographically laid out on the cortical surface and present a similar tuning across fractions and integers.

Note that, while previous studies have identified such tunings for non-symbolic numerosities presented as sets of dots (Harvey et al., 2013; Harvey & Dumoulin, 2017; Paul et al., 2022; Tsouli et al., 2021), symbolic tuning for number symbols has only been detected in occipito-temporal regions with functional magnetic resonance imaging (fMRI) (Cai et al., 2023), and in the mesial temporal lobe using intracranial recordings (Kutter et al., 2018, 2023, 2024). To maximize the chances of observing such tunings, we engaged participants in a demanding continuous number comparison task while collecting individual high-resolution (1.2 mm isotropic) fMRI data at high field (7 Tesla).

To anticipate our findings, we observed graded magnitude-based activation patterns in math-responsive areas as well as the medial temporal lobe. Within the math-responsive areas, numbers with closer magnitudes were represented by more similar activation vectors and in closer spatial proximity. In the bilateral anterior IPS / inferior parietal lobe (IPL), our images revealed a clear cortical semantic map with a shared magnitude code for both fractions and integers.

## Methods

### Participants

Twenty-two healthy adult participants were recruited for this study (13 women and 9 men, age range: 19-39 years, mean age = 26.4, SD = 5.6). All participants were right-handed, native French speakers, and held a bachelor’s degree in science. They had normal or corrected-to-normal vision and reported no history of psychiatric or neurological disorders. Written informed consent was obtained from all participants, who were compensated for their participation. The study was approved by the local ethics committee (Comité de Protection des Personnes Sud-Ouest et Outremer III; references: CEA 100 055, ID RCB 2020-A01713-36), and the study was conducted following the Declaration of Helsinki.

One participant was excluded due to a cyst in the temporal lobe that could affect that region’s anatomy and functional connectivity. Another participant was excluded after experiencing a panic attack in the scanner and requesting to leave it. Additionally, for a different participant, one of the response buttons failed, preventing analysis of his behavioral responses. As a result, we included 20 participants in the fMRI analyses and 19 in the behavioral analyses.

### Number comparison task

Participants were engaged in a serial continuous comparison task, inspired by Nelli et al. (2023), where each target had to be judged as larger or smaller than the previous one. The targets consisted of 12 numbers: six integers (i.e., 0, 1, 2, 3, 4, and 5), and six fractions 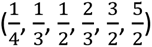. The stimuli were presented as white Arabic numerals on a black screen for 1 second, with fractions shown using a horizontal line. All transitions between the 12 target numbers occurred an equal number of times, with no number repeated consecutively, resulting in 11 presentations of each target and 132 unique transitions per run.

Before starting the fMRI experiment, participants completed a few trials to familiarize themselves with the task and stimuli. Each participant was assigned a number based on their arrival order. Participants with an even number were instructed to press the “left button” when the current number was greater than the previous one, and the “right button” when the current number was lower. In the second run, the buttons were reversed. Participants with an odd number started with the opposite button assignments. Each participant completed two runs, each lasting 9 min and 42 seconds. After the number presentation, a pseudo-randomly jittered interval of 2 to 4 seconds followed, fixed by the experimenter to ensure a mean SOA of 4 s, during which the fixation cross turned green if the response was recorded. Subsequently, the fixation cross turned red 500 ms before the onset of the next stimulus to prepare participants. Participants received a 10s rest time for every 45 stimuli, the fixation cross was on the screen the entire time, and the feedback with the percentage of correct answers appeared for 3s. Before starting the next block, the fixation cross turned red to prepare participants. At the beginning of each block, the last number before the rest was repeated when participants returned to the task.

### Math localizer

To localize math-responsive areas within each individual, we used a highly reproducible sentence-based paradigm previously employed multiple times in our lab (Amalric et al., 2018, 2023; Amalric & Dehaene, 2016, 2019, 2019; Pinel et al., 2007), which has the advantage of not including symbolic numbers in Arabic format. Participants read 60 sentences and had to decide whether they were True or False. Half of the sentences were mathematical statements, either stating arithmetic calculations in verbal form or describing geometrical facts concerning basic shapes (e.g., “*Number three multiplied by twelve is thirty-six*” or *“A rectangle has only one axis of symmetry”*). The remaining sentences were non-mathematical, including general factual knowledge or sentences referring to a person’s perspective (e.g., “*The camel is a large humpbacked mammal” or “For Einstein the atomic bomb is a harmless object”*). In each category, half of the sentences were true (T), and the other half were false (F), with the response buttons (left and right) counterbalanced across participants. Participants completed a single run lasting approximately 11 minutes.

The visual stimuli were balanced to control the number of words, total number of characters, word length, and frequency of occurrence in French. They were presented using rapid serial visual presentation (RSVP) at 350 ms per word, with an average total duration of 3.2 seconds. Each sentence was followed by a fixation cross, jittered to maintain an average stimulus onset asynchrony (SOA) of 8 seconds. A red cross was presented 350 ms in advance of the upcoming stimulus. Participants were instructed to answer as quickly and as accurately as possible, and response times were measured relative to the onset of the last word.

Stimulus delivery and response collection of both Localizer and Numbers tasks were controlled by Expyriment (Krause & Lindemann, 2014).

### fMRI acquisition

MRI data were acquired using a 7T MAGNETOM whole-body MR scanner (Siemens Healthineers, Erlangen, Germany) with a 32-channel head coil (Nova Medical, Wilmington, USA) at the NeuroSpin Center of the French Alternative Energies and Atomic Energy Commission. Structural MRI data were collected using T1-weighted rapid gradient echo (MPRAGE) sequence (repetition time (TR) = 5000 ms, echo time (TE) = 2.51 ms, resolution = 0.65 mm isotropic, field of view (FoV) = 208⨯208 mm2, flip angle = 5/3, bandwidth (BW) = 250 Hz/px, echo spacing = 7 ms). Functional MRI (fMRI) data were acquired using a T2*- weighted gradient echo planar imaging (EPI) sequence (TR =2000 ms, TE = 21 ms, voxel size = 1.2 mm isotropic, multiband acceleration factor = 2, FoV = 192⨯192 mm2, flip angle=75, BW=1488 Hz/px, partial Fourier = 6/8, echo spacing = 0.78 ms, number of slices = 70 (no gap)). To correct for EPI distortion, a five-volume functional run with the same parameters except for the opposite phase encoding direction (posterior to anterior) was acquired immediately before each task run. Manual interactive shimming of the B0 field was performed for all participants. The system voltage was set at 250 V for all sessions, and the fat suppression was decreased per run to ensure that the specific absorption rate did not surpass 62% for all functional runs.

### Anatomical and Functional Preprocessing

Both anatomical and functional data were preprocessed using fMRIPrep 20.0.2. This pipeline includes standard preprocessing steps organized in an easy-to-use workflow that ensures robustness independently of the data idiosyncrasies and high consistency of results (Esteban et al., 2019). The T1-weighted (T1w) image was corrected for intensity non-uniformity (INU) with N4BiasFieldCorrection (Tustison et al., 2010), distributed with ANTs 2.3.3 (Avants et al., 2008), RRID:SCR_004757), and used as T1w-reference throughout the workflow. The T1w-reference was then skull-stripped with a *Nipype* implementation of the antsBrainExtraction.sh workflow (from ANTs), using OASIS30ANTs as target template. Brain tissue segmentation of cerebrospinal fluid (CSF), white-matter (WM) and gray-matter (GM) was performed on the brain-extracted T1w using fast (FSL 6.0.5.1:57b01774, (Zhang et al., 2001). Brain surfaces were reconstructed using recon-all (FreeSurfer 7.2.0, (Dale et al., 1999), and the brain mask estimated previously was refined with a custom variation of the method to reconcile ANTs-derived and FreeSurfer-derived segmentations of the cortical gray-matter of Mindboggle (Klein et al., 2017). Volume-based spatial normalization to one standard space (MNI152NLin2009cAsym) was performed through nonlinear registration with antsRegistration (ANTs 2.3.3), using brain-extracted versions of both T1w reference and the T1w template.

A reference volume and its skull-stripped version were generated by aligning and averaging 1 single-band reference (SBRefs). Head-motion parameters with respect to the BOLD reference (transformation matrices, and six corresponding rotation and translation parameters) are estimated before any spatiotemporal filtering using mcflirt (FSL 6.0.5.1:57b01774, (Jenkinson et al., 2002). The estimated *fieldmap* was then aligned with rigid registration to the target EPI (echo-planar imaging) reference run. The field coefficients were mapped onto the reference EPI using the transform. BOLD runs were slice-time corrected to 0.975s (0.5 of slice acquisition range 0s-1.95s) using 3dTshift from AFNI (Cox & Hyde, 1997). The BOLD reference was then co-registered to the T1w reference using bbregister (FreeSurfer) which implements boundary-based registration (Greve & Fischl, 2009). Co-registration was configured with nine degrees of freedom to account for distortions remaining in the BOLD reference. First, a reference volume and its skull-stripped version were generated using a custom methodology of *fMRIPrep*. Several confounding time series were calculated based on the *preprocessed BOLD*: framewise displacement (FD), DVARS, and three region-wise global signals. FD was computed using two formulations following Power (absolute sum of relative motions, (Power et al., 2014) and Jenkinson (relative root mean square displacement between affines, (Jenkinson et al., 2002). FD and DVARS are calculated for each functional run, both using their implementations in *Nipype* (following the definitions by (Power et al., 2014). The three global signals are extracted within the CSF, the WM, and the whole-brain masks. Additionally, a set of physiological regressors was extracted to allow for component-based noise correction (*CompCor*, (Behzadi et al., 2007)). Principal components are estimated after high pass filtering the *preprocessed BOLD* time series (using a discrete cosine filter with 128s cut-off) for the two *CompCor* variants: temporal (tCompCor) and anatomical (aCompCor). tCompCor components are then calculated from the top 2% variable voxels within the brain mask. For aCompCor, three probabilistic masks (CSF, WM, and combined CSF+WM) are generated in anatomical space. The implementation differs from that of (Behzadi et al., 2007) in that instead of eroding the masks by 2 pixels on BOLD space, a mask of pixels that likely contain a volume fraction of GM is subtracted from the aCompCor masks. This mask is obtained by dilating a GM mask extracted from the FreeSurfer’s *aseg* segmentation, and it ensures components are not extracted from voxels containing a minimal fraction of GM. Finally, these masks are resampled into BOLD space and binarized by thresholding at 0.99 (as in the original implementation). Components are also calculated separately within the WM and CSF masks. For each CompCor decomposition, the *k* components with the largest singular values are retained, such that the retained components’ time series are sufficient to explain 50 percent of variance across the nuisance mask (CSF, WM, combined, or temporal). The remaining components are dropped from consideration. The head-motion estimates calculated in the correction step were also placed within the corresponding confounds file. The confound time series derived from head motion estimates and global signals were expanded with the inclusion of temporal derivatives and quadratic terms for each (Satterthwaite et al., 2013). Frames that exceeded a threshold of 0.5 mm FD or 1.5 standardized DVARS were annotated as motion outliers. Additional nuisance time series are calculated by means of principal components analysis of the signal found within a thin band (*crown*) of voxels around the edge of the brain, as proposed by (Patriat et al., 2017). The BOLD time series were resampled into standard space, generating a *preprocessed BOLD run in MNI152NLin2009cAsym space*. First, a reference volume and its skull-stripped version were generated using a custom methodology of *fMRIPrep*. All resamplings can be performed with *a single interpolation step* by composing all the pertinent transformations (i.e., head-motion transform matrices, susceptibility distortion correction when available, and co-registrations to anatomical and output spaces). Gridded (volumetric) resamplings were performed using antsApplyTransforms (ANTs), configured with Lanczos interpolation to minimize the smoothing effects of other kernels (Lanzcos, 1964). Non-gridded (surface) resamplings were performed using mri_vol2surf (FreeSurfer).

### Data analysis

We analyzed behavioral data by plotting the efficiency score (Accuracy/RTs) for each combination of the 12 numbers and as a function of the numerical distance between the current and previous number.

Statistical analyses of the fMRI data were performed using NiLearn (RRID: SCR_001362) and rsatoolbox (van den Bosch et al., 2025). In the general linear models (GLM), regressors were convolved with the canonical hemodynamic response function (HRF). GLM1 included 17 regressors of interest: one for each current target number (12 regressors); two for left and right button presses; one for RT difficulty, calculated as the current RT minus the participant’s average RT; one for feedback onset; and one for the first target presentation, where participants were not required to respond. Additionally, to regress out motion and physiological noise, we included six head motion parameters, the CSF and white matter signal, 10 anatomical CompCor (computed by *fMRIPrep*), and 14 cosine drifts. This GLM was used for the PCA, RSA, searchlight, MDS, and Gaussian fit analyses.

The data remained unsmoothed for all analyses, but for visualization purposes, we smoothed the data at the group level with a 6 mm kernel. An exception was made for the Gaussian fit analysis, where the data were smoothed at the individual level with a 4 mm kernel, but no smoothing was applied at the group level.

GLM2 was very similar to GLM1, except that we excluded the RT difficulty regressor and included the logarithm of the absolute distance, calculated as log(|current number - previous number|). This model was used exclusively for Figure 3D, and the data were smoothed at the group level with a 6 mm kernel.

### Principal Component Analysis (PCA)

We used PCA on the 12 target-related voxel-wise model weights to identify the most important orthogonal dimensions for the task. We averaged the t-maps for each target number across participants using unsmoothed data (similar results were obtained at the individual level in the majority of participants). Then, we created a matrix where each row corresponded to a target number, and each column corresponded to a voxel. We applied PCA to these weights. We determined that two components were optimal by examining the elbow of the Scree Plot. To project the results onto the cortical surface of a common space across participants, we selected the top 1% of voxels, and this data was subsequently smoothed with a 6 mm kernel.

### Representation Similarity Analysis (RSA)

We identified brain areas showing greater activation for math-related sentences compared to non-math-related sentences in the localizer task, at the whole brain level for each participant (each thresholded at p < 0.001 uncorrected). From the contrast math > non-math sentences, we also manually selected the parietal, frontal, and temporal ROIs at the individual level. BOLD representation dissimilarity matrices (RDMs) were constructed by calculating the correlation distance d=1-r, where r is the Pearson correlation coefficient between the within-subject multivoxel patterns elicited by each number, resulting in a 12×12 RDM for each participant, which was then averaged across participants. To visualise neural geometry, we used a two-dimensional metric inter-individual difference scaling (INDSCAL) MDS model, which takes into account the individual participants’ RDMs, implemented using the R-package SMACOF (Flesch et al., 2022; Leeuw & Mair, 2009).

To identify cortical sectors in which the neural RDM correlated with a specific RDM – particularly focusing on the distance across the domains of fractions and integers – while accounting for the influence of remaining RDMs, we performed a multiple-regression searchlight RSA. This analysis was performed within a mask of mathematically responsive areas defined by the localizer and used in the RSA (masking the union of voxels activated to math > non-math sentences in one or more participants), along with Glasser-defined ROIs of hippocampi and parahippocampal cortex. The neural RDM for each searchlight was computed using correlation distances. For the searchlight analysis, we included all voxels within a radius of 4mm of the central voxel. For each searchlight sphere, we computed neural RDMs from the condition-by-voxel matrix of estimated neural responses using Pearson correlation distance between pairs of conditions. This procedure was carried out at the individual participant level, followed by a second-level analysis across participants, using FDR correction q < 0.001 for multiple comparisons. The multiple regression model included five predictor RDMs: distance within fractions, category of fractions, distance within integers, category of integers, and distance across the domains of integers and fractions.

### Voxel tuning

For this analysis, we averaged the t-maps for each target number across participants using smoothed data (4 mm FWHM). The same analysis was also performed at the individual level in each participant’s native space. We then created a matrix in which each column represented a target t-value and each row corresponded to a voxel within the mask of math-related regions defined by the localizer math > non-math and used in the RSA, as well as the Glasser ROIs of parahippocampal, and hippocampus. Separate analyses were conducted for integers and fractions. Each voxel’s values were normalized between their minimum and maximum across conditions of integers or fractions, resulting in a range from 0 to 1. A Gaussian function was then fitted using the Levenberg-Marquardt method within a bounded parameter space, yielding for each voxel a preferred value on a continuous scale from 0 to 5 for integers and from 0.25 to 2.5 for fractions. The amplitude was constrained between 0 to 1, and the width between 0.1 to 2. Only voxels with a goodness of fit (R^2^) greater than 0.5 were included, and at the individual level we retained the voxels with R^2^ >0.3. Using data from one of the runs (1 or 2), we categorized voxels into 6 bins by rounding their preferred values for integers and, for fractions, by applying category boundaries based on the subtraction (next fraction - previous fraction)/2. Then, we plotted the normalized activation using data from the other run and averaged the two resulting plots (finding preferred numbers with run 1 and plotting the activation of run 2, and vice versa). To assess statistical significance, we organized the average data in a matrix (6×6 for integers, 6×6 for fractions) where columns indicated the preferred numbers and rows the average normalized activation to each target number. Then, we tested whether the diagonal elements of this matrix were significantly larger than the off-diagonal elements. For that, we shuffled the columns to break the correspondence between rows and columns and computed a permutation test (n = 720 permutations).

We correlated each voxel’s preferred value for fractions and for integers within math-responsive areas and in aIPS/IPL (Glasser ROIs: PF and IP2). To assess the significance of these correlations, we conducted a permutation test by randomly shuffling one of the vectors 10000 times and comparing the observed Pearson correlation with the distribution obtained from the shuffled data. This allowed us to test whether the observed correlation between fractions and integers was higher than expected by chance. We then applied the same procedure to assess the correlation between the two runs for integers and fractions. To plot the tuning data across fractions and integers, we selected the voxels corresponding to numbers from 0 to 3, as there were too few voxels representing magnitudes of 4 and 5. Voxels were assigned to each category based on their rounded preferred magnitude (e.g., voxels categorized as 0 had preferred values between 0 and 0.49). We then extracted the normalized activation for fractions from these voxels (see Supplementary Materials Figure S2).

## Results

### Behavioral evidence for a distance effect and a mental number line

Participants responded significantly more slowly to fractions (MFrac-Frac = 1192 ms; SDFrac-Frac = 129) than to integers (MInt-Int = 808 ms; SDInt-Int = 64; *t*(28) = −17.12, *p* < 0.001), and they made more errors when both the previous and current numbers were fractions (MFrac-Frac = 9.5%; SDFrac-Frac = 6.90) than when both were integers (MInt-Int = 3.7%, SD Int-Int = 3.56; *Mann-Whitney U = 368*, z = 3.77, *p* < 0.001). In trials requiring comparisons between integers and fractions, participants were also slower when integers preceded fractions (MInt-Frac = 1053 ms; SDInt-Frac = 191) than when fractions preceded integers (MFrac-Int= 952 ms; SDFrac-Int = 147, *Mann-Whitney U = 469*, z = 2.14, *p* = 0.03), with no corresponding differences in accuracy between the two conditions (MInt-Frac = 8.11 % of errors; SDInt-Frac = 8.11; MFrac-Int = 5.56 % of errors; SDFrac-Int = 5.40; *Mann-Whitney U = 329*, z = 1.23, *p* = 0.22). These results were stable across time, since reaction times were highly correlated between run 1 and run 2 (*r* = 0.95, *p* < 0.0001), as were the percentage of errors (*rho* = 0.49, *p* < 0.0001). Overall, these results indicate that participants found conditions involving fractions more difficult than those involving integers.

RTs and the percentage of errors are presented as a function of absolute distance in Fig. 1B and C for each condition (i.e., Frac-Frac, Frac-Int, Int-Frac, Int-Int). Regardless of condition, we found a statistically significant distance effect: participants were slower and less accurate when two numbers were closer on the number line. This effect was particularly pronounced when both the current and previous numbers were fractions (RTs: βFrac-Frac = −220.26, *t*(28) = −6.54, *p* < 0.0001; Errors: βFrac-Frac = −11.54, *t*(28) = −6.26, *p* < 0.0001; βs obtained with a linear regression model with log-transformed numerical distance as predictor), but a distance effect was also present when both numbers were integers (RTs: βInt-Int = −146.68, *t*(28) = −3.60, *p* = 0.001; Errors: βInt-Int = −6.65, *t*(28) = −2.75, *p* = 0.01) or when participants compared an integer to a previous fraction (RTs: βFrac-Int = −252.01, *t*(34) = −5.12, *p* < 0.0001; Errors: βFrac-Int = −9.69, *t*(34) = −5.54, *p* < 0.0001) and vice-versa (RTs: βInt-Frac = −308.79, *t*(34) = −4.61, *p* < 0.0001; Errors: βInt-Frac = −12.67, *t*(34) = −4.37, *p* < 0.0001).

**Figure 1.**
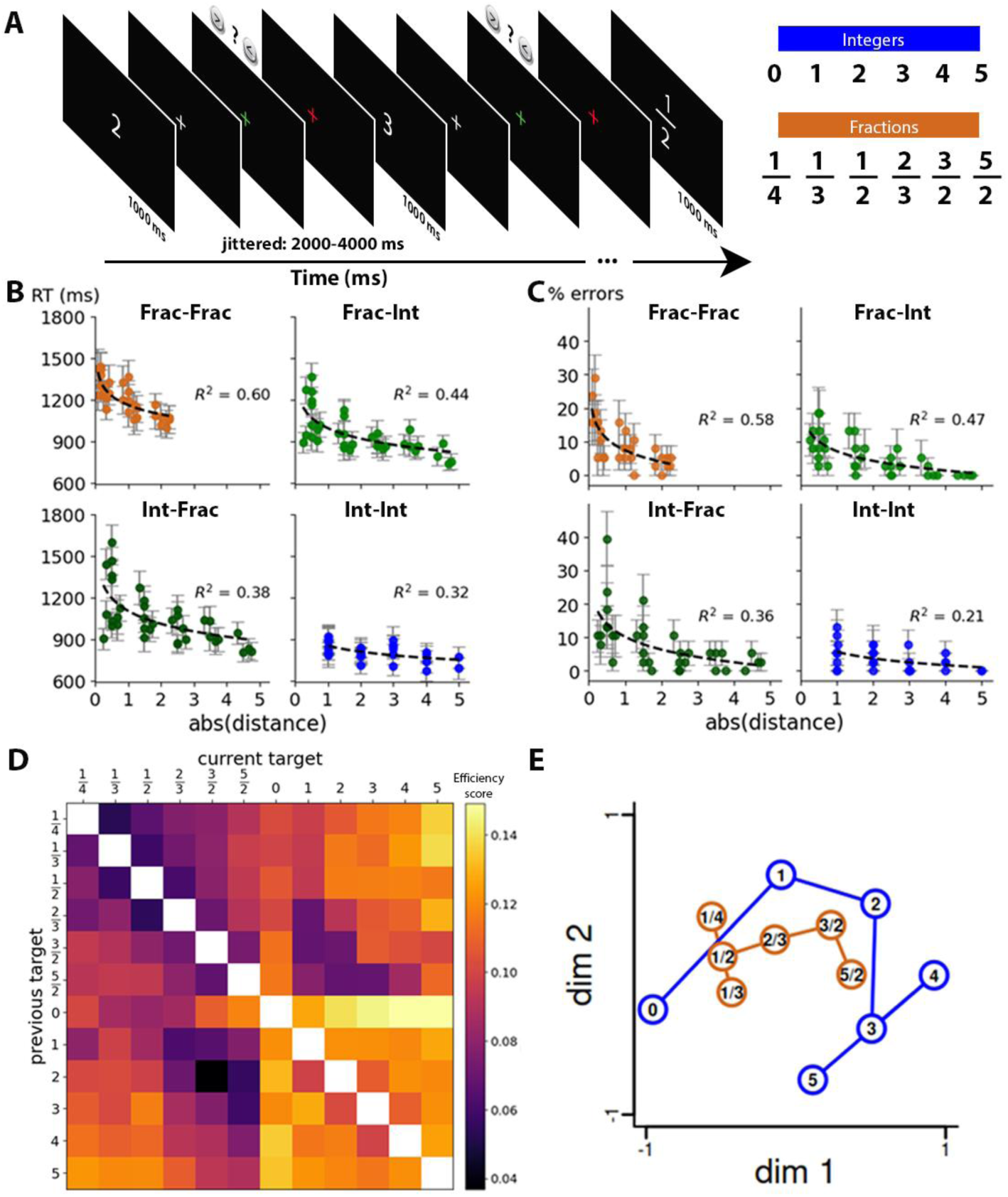
Experimental design and behavioral results. **A)** An example of the task sequence in the scanner: participants compared the current number to the previous one. Each number was followed by a fixation cross that turned green when the response was registered and red 500 ms before the next trial to prepare participants. After 45 stimuli, participants received feedback on their percentage of correct answers. Right of panel A, the full set of stimuli is shown, including both integers and fractions. **B)** RTs as a function of absolute distance between current and previous numbers (132 combinations). Panels separate trials in which both the previous and the current numbers were integers (int), both were fractions (frac), and trials with mixed number domains. **C)** Percentage of errors as a function of the absolute distance between current and previous numbers (same format). **D)** Efficiency Score, where smaller values (closer to purple) indicate greater difficulty in comparing numbers, while larger values (yellow) indicate easier comparisons. **E)** Two-dimensional MDS representation of the efficiency score matrix, showing that fractions and integers are organized according to their magnitudes.

To combine information about RTs and the percentage of errors, we calculated the efficiency score (Accuracy / RTs) for each pair of numbers (see Fig. 1D). A multiple linear regression revealed a logarithmic distance effect between the previous and current numbers (β = 0.03, *t*(7) = 3.55, *p* = 0.0006), a lesser efficiency when the previous item was a fraction compared to an integer (β = 0.015, *t*(7) = 3.76, *p* = 0.0003), and also when the current item was a fraction compared to an integer (β = 0.026, *t*(7) = 6.64, *p* < 0.0001). The only statistically significant interaction was between the previous and current number conditions, indicating further difficulty when both items were fractions (β = 0.011, *t*(7) = 2.17, *p* = 0.03). The model explained a substantial proportion of the variance (R² = 0.74).

MDS of the efficiency score (Figure 1E) showed an approximately one-dimensional curvilinear structure (number line) embedded in two dimensions, similar to that previous observed in similar magnitude comparison tasks (Nelli et al., 2023; Sheahan et al., 2021). Importantly, fractions and integers were integrated together, and both were ordered according to their magnitudes on the first dimension (x axis), with smaller numbers on the left side and larger numbers on the right side. The second dimension (y axis) seemed to encode difficulty, as it separated the intrinsically easier numbers 0 and 5 (because they lie at the extremes) from other numbers. After three, the number line was bent into a zig-zag pattern, which might occur because these numbers were easier, as fractions only spanned the magnitude of integers up to 3. When we recomputed the MDS excluding 0 and 5, their number line organization was largely unchanged (Figure S1), further supporting the role of magnitude as a major organizing dimension of semantic space.

### Univariate fMRI activations converge to a classic network of math-responsive areas

Three univariate contrasts converged to a largely common set of classic math-responsive regions (Figure 2). First, a contrast for greater activation to math-related sentences than to non-math-related sentences in the localizer revealed an extensive bilateral network, including the full length of the IPS, posterior inferior temporal gyri (pITG), precentral gyri, middle frontal gyri (MFG), and precuneus. The opposite contrast (in blue), indicating a greater response to non-math-related sentences, activated a classic and bilateral network of semantic regions, comprising the angular gyri, superior and middle temporal gyri, temporal poles, fusiform gyri, superior and inferior frontal gyri, and posterior cingulate. Second, in the numbers task, most of the above math-responsive network showed a greater activation for fractions compared to integers in bilateral regions, including the IPS, precentral gyri, inferior and middle frontal gyri, precuneus, pITG, now also extending to lateral fusiform and lateral occipital cortex (presumably due to the higher visual complexity of fractions). Third, similar regions, including bilateral IPS and pITG showed a distance effect during number comparison, i.e. a parametric modulation by the log of the absolute distance between the current and previous numbers, with greater activation to smaller distances, as previously described (e.g. Pinel et al., 2001).

**Figure 2:**
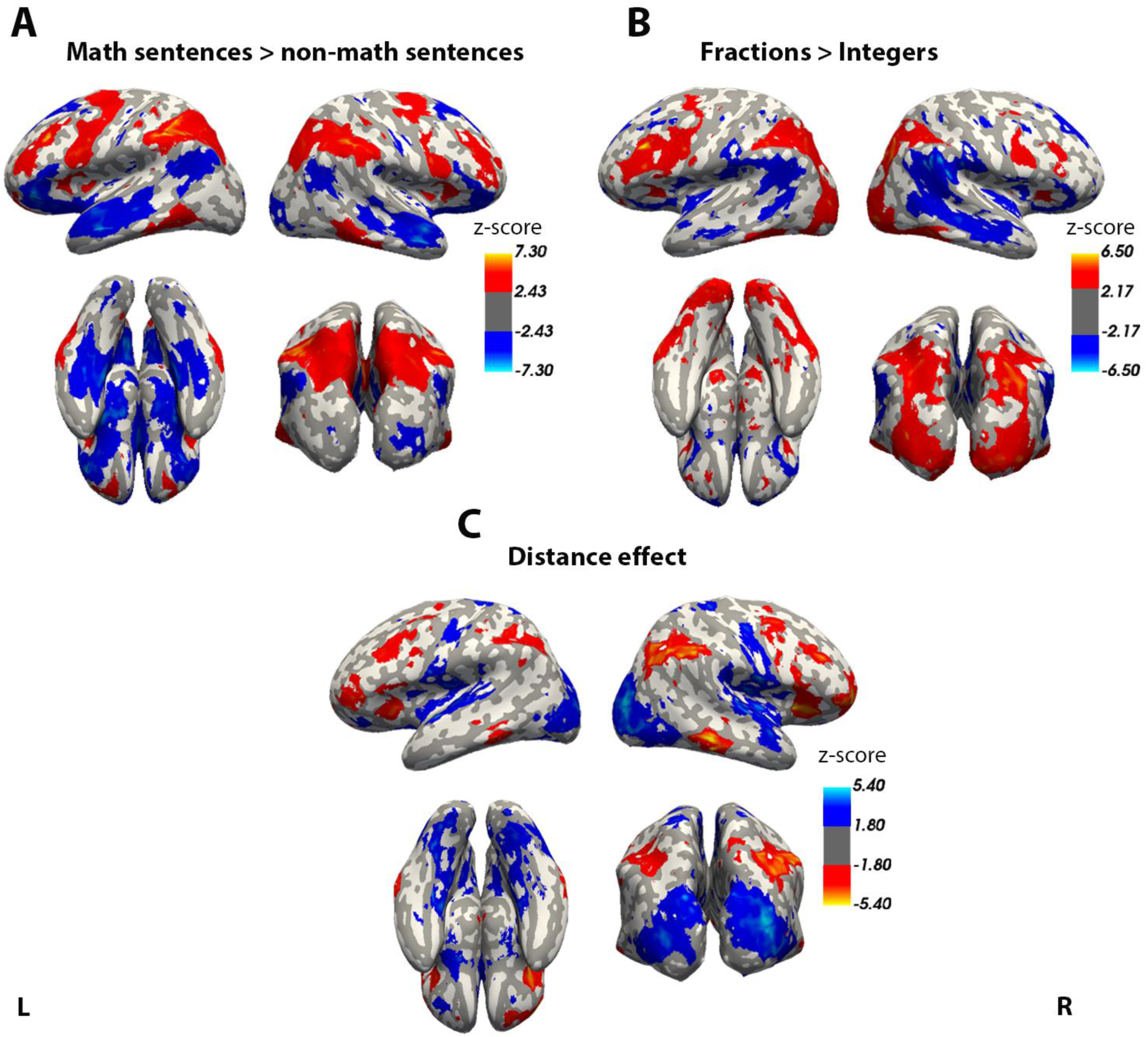
A math-responsive cortical network and its response to three distinct univariate contrasts. **A)** Greater activation to math than to non-math sentences in the localizer. **B)** Greater activation to fractions than to integers. **C)** Effect of the distance between the current and previous numbers (greater activation to closer numbers). All maps thresholded at false detection rate (FDR) q< 0.05 across the whole brain.

### Vector-based analyses of the neural representation of numbers

Using a GLM with one regressor per target, we obtained, within each participant, a reliable high-resolution estimate of the fMRI activation pattern or “vector” evoked by each of 12 individual number concepts. Our aim was to probe whether these vectors organized as a semantic map and what were its main axes of variations.

As a first step, we performed a whole-brain principal component analysis (PCA) of the average vectors across participants within a standard anatomical space (MNI). PCA revealed two main orthogonal components that explained most of the variance in our data: PC1 reflected a categorical distinction between fractions and integers, while PC2 reflected a near-perfect ordering by number magnitude, from small to large, distinguishing smaller and larger numbers while pooling together fractions and integers (Figure 3). In individual analyses, the first component was observed in 14 of our 20 participants, whereas the second was present in 13. Each component produced distinct spatial maps, although some parietal regions were involved in more than one component. Specifically, PC1 confirmed the univariate analysis of Figure 2B, indicated that fractions (probably because of their greater visual and/or semantic complexity), mobilized bilateral occipital, pITG, posterior IPS and MFG. Crucially, PC2 revealed that a magnitude map mixing fractions and integers was present in much of the math-responsive network (Figure 3, bottom), with a particularly strong small-to-large gradient in the anterior IPS (extending in to the inferior parietal lobule for small numbers), with additional contributions of the MFG, pITG, posterior cingulate, parahippocampal cortex and hippocampus. Together, PC1 and PC2 explained 36.23% of the variance. Thus, this analysis confirms that number magnitude is a major determinant of number representations which affects a broad set of brain areas.

**Figure 3.**
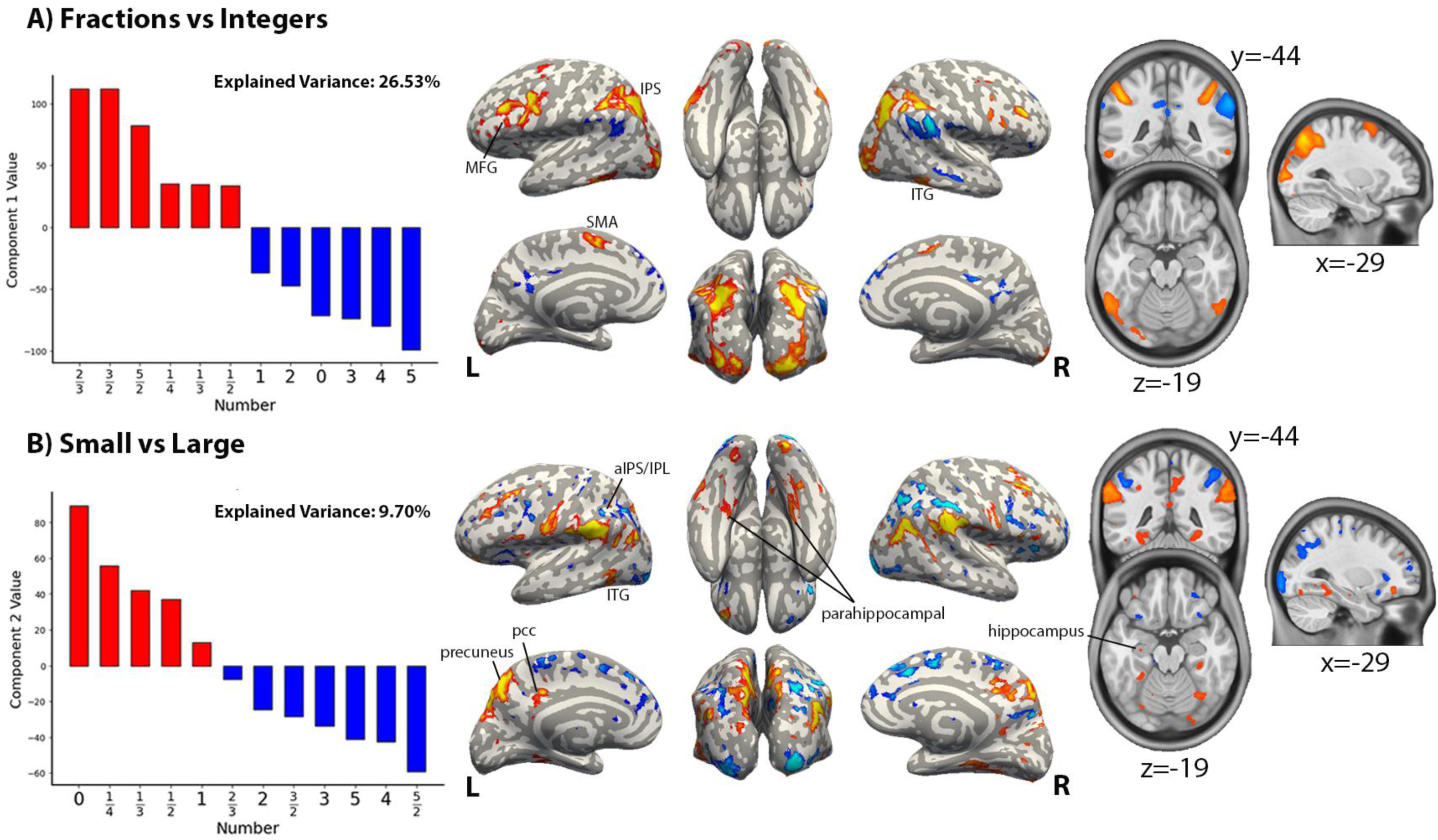
Whole-brain principal Component Analysis (PCA) reveals the two main dimensions of number representations: category and magnitude. In this purely data-driven analysis of the activation vectors evoked by each of the 12 number target, PC1 (top) distinguished between fractions and integers, while PC2 order numbers almost perfectly according to their magnitude. In each case, the 12 loadings (ordered) and the corresponding voxel weights are shown, in surface and slice views. **SFG:** superior frontal gyrus; **MFG:** middle frontal gyrus; **IFG:** inferior frontal gyrus; **SMA:** supplementary motor area; **PCC:** posterior cingulate cortex; **ITG:** inferior temporal gyrus; **MTG:** middle temporal gyrus

To take advantage of the high-resolution individual activations available using 7T fMRI, we next extracted single-participant vectors within the entire math-responsive network by first identifying subject-specific voxels showing greater activation to math than to non-math sentences in the localizer (p < 0.001 uncorrected). Then, we focused on the six main areas identified by Amalric et al. (2016) (left and right IPS, pITG, and MFG). In those voxels, we extracted the activation vectors to each of the 12 target numbers in the main comparison task, then visualized those vector spaces using RDMs followed by MDS.

This analysis confirmed that neural vector spaces were organized by two dimensions: a main distinction between fractions and integers, and a gradient of continuous magnitude (Figure 4). The latter was seen in two ways: in the RDMs, neural vectors became increasingly similar as the numbers were closer, i.e., for cells closer to the diagonal of the matrix (for both integers and fractions); and in the 2-dimensional MDS solutions of those RDMs, integers and fractions, while separated, were similarly ordered by magnitude. Furthermore, information was represented similarly across the three math-responsive regions IPS, pITG and MFG.

**Figure 4.**
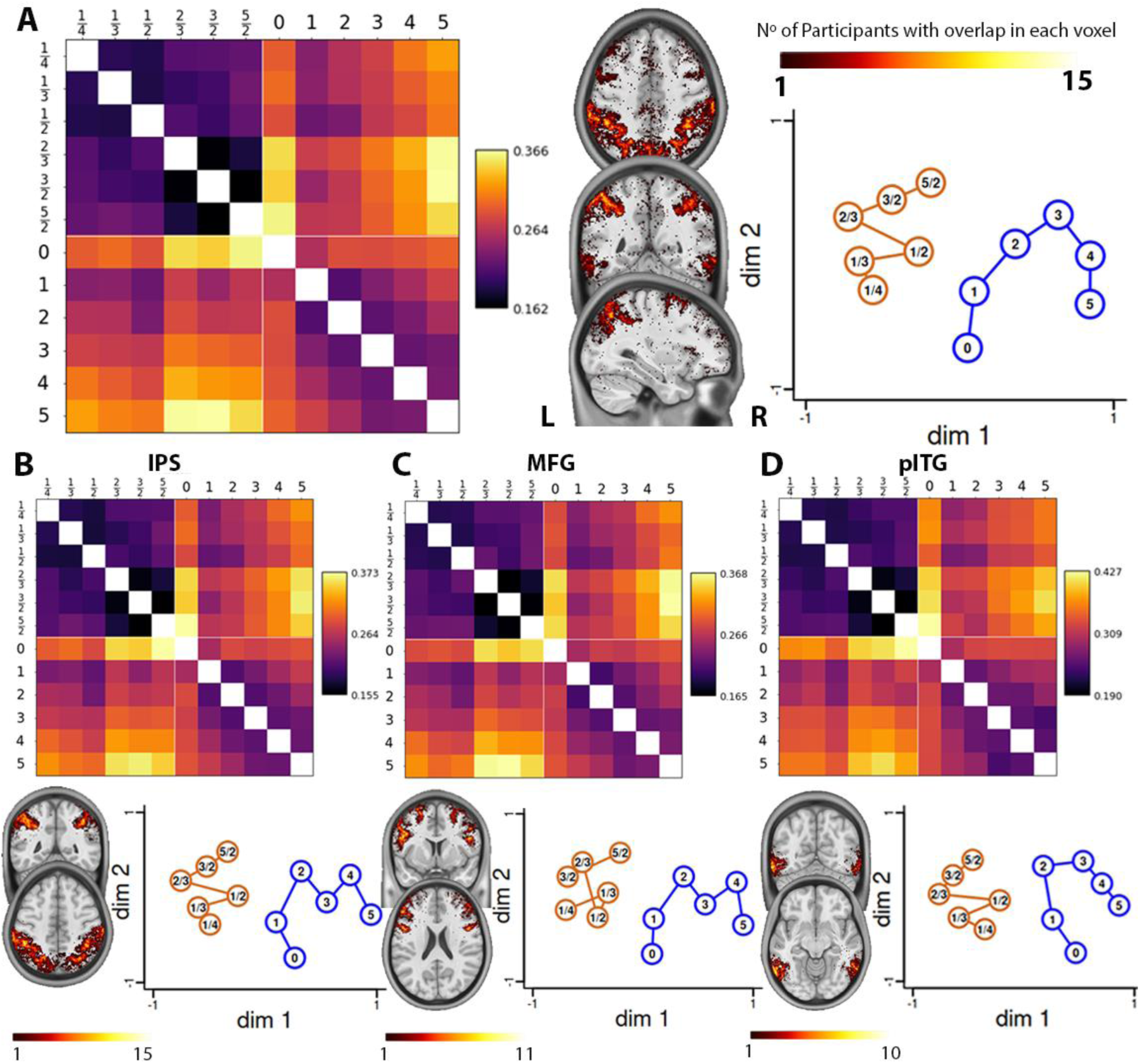
Visualization of the neural representational spaces for numbers within math-responsive areas. Representational similarity matrices and their 2-dimensional MDS solutions were used to visualize the 12-dimensional vector spaced spanned by number activations within individual voxels-of-interest identified by the math>non-math contrast in the sentence localizer. **A)** Activations pooled On the left, the RDM shows the average dissimilarity across participants. In the middle, voxel overlap across individuals is displayed with lighter colors indicating higher overlap (up to a maximum of 15 participants). On the right is the corresponding neural geometry representation derived from MDS. **B)** The top panel shows the average RDM restricted to parietal regions of interest (ROIs); the bottom panel shows participant overlap and the resulting neural geometry. **C)** RDM and MDS for frontal ROIs. **D)** RDM and MDS for temporal ROIs.

**Figure 5.**
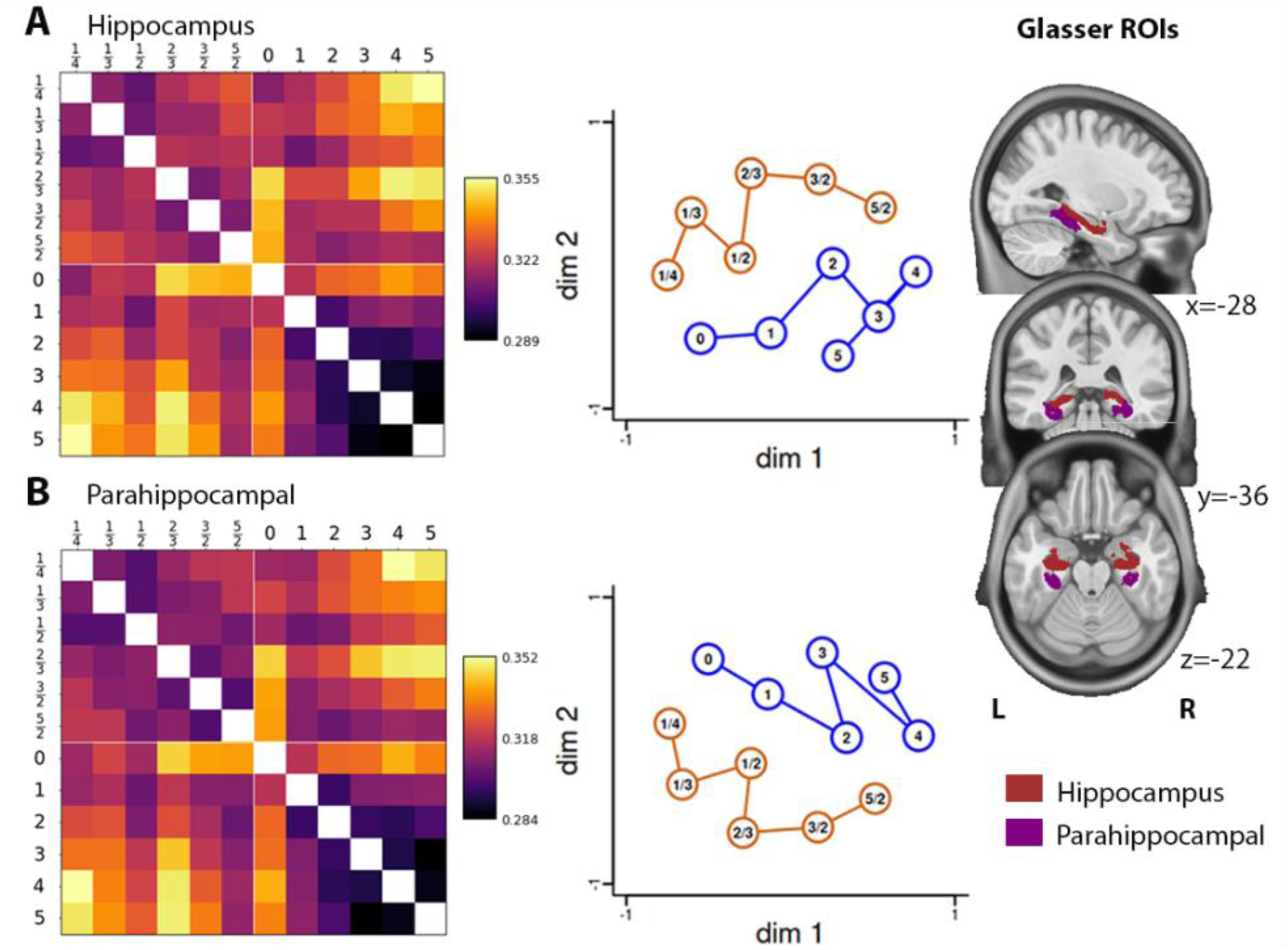
Number line representations within the medial temporal lobe. Same format as figure 4. A, hippocampus, B, parahippocampal regions. Both regions were taken from the Glasser atlas and applied to individual participants.

Since the hippocampus and parahippocampal cortex were also identified in the PCA, and since the human hippocampus and mesial temporal lobe have been shown to contain number-tuned neurons (Kutter et al., 2018, 2023, 2024), we also analyzed vector spaces in those regions. We used the bilateral ROIs defined by Glasser et al., 2016 and extracted the entire hippocampus and parahippocampal cortex. In these regions, an organization according to magnitude was even more prominent, with MDS dimension 1 ordering the targets according to their approximate magnitude regardless of whether they were fractions or integers (although a distinction between the two domains remained visible in MDS dimension 2). Note that, similar to the behavioral data, the integer number line seemed to bend after three in all MDS representations, which may reflect the absence of fractions in this range. Importantly, this effect was not driven solely by the presence of “easy” extreme numbers, as the pattern remained consistent when we recomputed the MDS excluding the extremes 0 and 5 (Figure S1).

To explicitly test for a shared representation of fractions and integers, we modelled the RDMs using multiple regression analysis focusing on the distance between fractions and integers, but also including 4 regressors controlling for categorical effects and for within-domain distance effects (Figure 6, top). In each region used in RSA, a significant effect of fraction-to-integer distance on neural dissimilarity was found (Table S1). Thus, math-responsive regions were not only organized according to within-domain distances between two integers or two fractions (e.g., 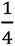 is more similar to 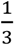 than to 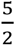), but also by the between-domain distances (e.g., 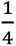 is more similar to 0 than to 4), indicating that in our educated adult participant, the magnitude of fractions was well integrated with that of integers. A searchlight RSA analysis within the math-responsive areas (Figure 6, bottom) further showed that the regions sensitive to cross-domain distances included the bilateral hippocampus and parahippocampal cortex, IPS, pITG, MFG as well as precentral cortex, posterior cingulate cortex and precuneus.

**Figure 6.**
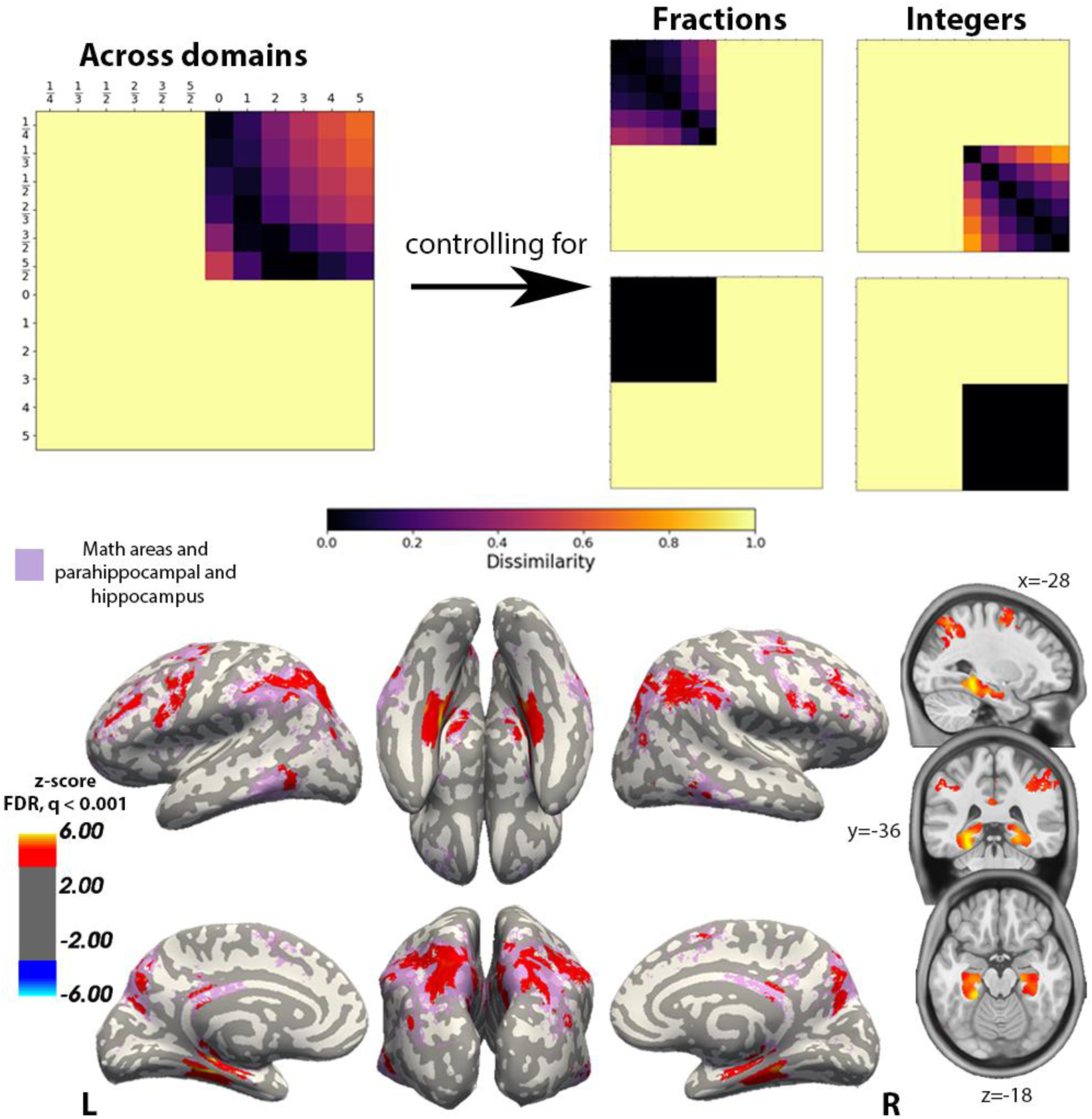
Brain areas sensitive to the numerical distance between fractions and integers. Top, theoretical RDMs used to model the empirical RDMs. Our focus was on the upper-right quadrant (numerical distance *between* integers and fractions) while controlling for categorical and distance effects *within* each domain. A whole-brain searchlight analysis revealed that regions encoding distance between domains (in red) included the hippocampus and parahippocampal cortex, as well as cortical regions of the IPS, pITG, MFG and precentral cortex, precuneus, and posterior cingulate, overlapping strongly with math-responsive areas identified by the sentence localizer (in purple).

### Individual voxel tuning and numerotopic map

The similarities observed so far between neural activation vectors could, in principle, arise either from distributed representations or from sparse localized neural responses. Furthermore, in the latter case, cortical patches responding to different concepts could either be distributed randomly on the cortical surface or form an overall mesoscopic topographic map (Deniz et al., 2019; Huth et al., 2016). Indeed, in the number domain, previous research showed that the firing of individual neurons can be fitted by a Gaussian tuning curve, reflecting their preference for a specific quantity (Kutter et al., 2018, 2023, 2024; Nieder & Dehaene, 2009). Furthermore, those preferences cluster on the cortical surface to form “numerotopic” maps that are detectable by fMRI (Cai et al., 2023; Harvey et al., 2013; Harvey & Dumoulin, 2017). Inspired by this prior work, we fitted the present single-voxel fMRI data with Gaussian tuning curves in order to characterize how numerical information is spatially represented and whether a spatial map exists in which voxels corresponding to similar magnitudes are located closer together, regardless of domain (integers or fractions).

We extracted the single-voxel group-average responses to all numbers within the math-responsive areas defined by the localizer. Each voxel’s response profile was fitted by a Gaussian, separately for integers and fractions and for each run, keeping only the voxels that showed greater than 50% variance explained. The presence of reliable voxel tunings was indicated by the fact that, across voxels, the numerical preferences computed in run 1 and run 2 were significantly and positively correlated (r = 0.39, *p* = 0.0001 for integers, and r = 0.59, *p* = 0.0001 for fractions; Figure 7A). There was also a smaller but still reliable correlation between preferred fraction and preferred integer (r = 0.01, *p* = 0.008). For brain visualization, in Fig. 7A we used the two runs. Figure 7B shows the average cross-validated voxel response profiles, after pooling across all the aforementioned voxels (voxel preferences were determined by fitting the data from one run, and then the normalized activation was plotted from the other run). A statistical test confirmed that the preferred number identified in one run indeed led to significantly stronger activation to that target number than to others in the other run (for integers *p* = 0.003, and for fractions *p* = 0.006).

**Figure 7.**
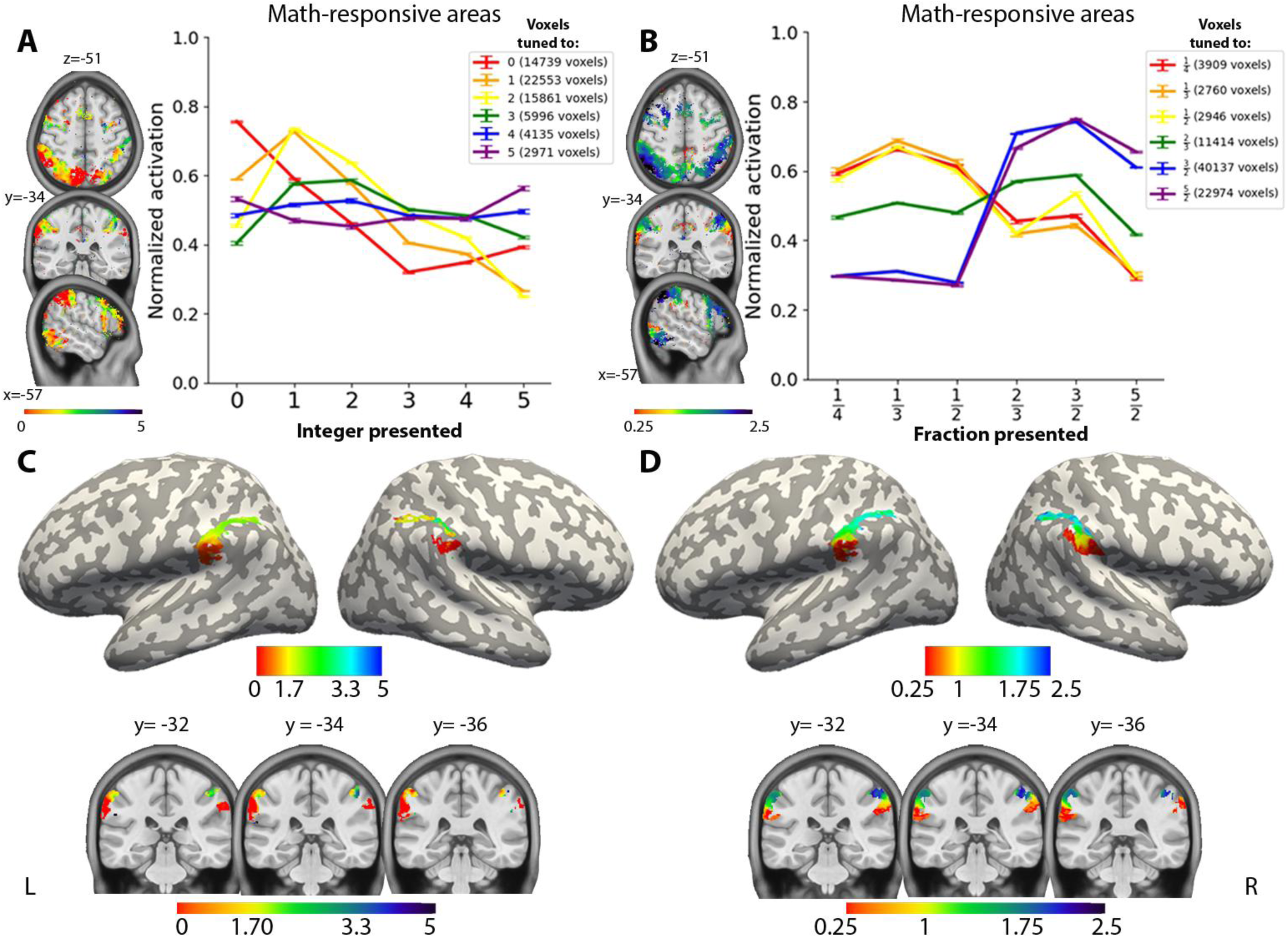
Evidence of voxel tuning to specific numbers and of an overall numerotopic map. A) Correlation, across voxels, between the preferred numbers identified in two independent data sets (run 1 and run 2). B) Cross-validated plots of normalized activations within voxels identified by Gaussian fitting as tuned to a specific integer (top) or a specific fraction (bottom). All voxels within math-responsive areas were included in this plot. C) Surface view (left) and axial slices (right) showing voxel-preferences for fractions (top) and integers (bottom) across several voxels within anterior IPS and IPL. This bilateral region comprises a congruent numerotopic map across integers and fractions, in which larger numbers are represented with bluish hues and smaller numbers with reddish hues. The map is shown using data averaged across participants; see Figure S1 for examples single-subject maps.

For integers, a clear distance effect can be seen in Figure 7B, with activation decreasing as the numerical distance to the preferred number increased. This effect was prominent for voxels tuned to numbers 0, 1, 2, and to a lesser extent 3, but became undetectable for numbers 4 and 5, probably because very few voxels were tuned to the latter numbers. For fractions, the distance effect was less evident, perhaps because magnitudes were closer, but a reliable distinction was observed between voxels tuned to fractions 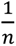, those tuned to fraction 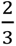, and those tuned to fractions larger than 1. No such reliable tuning was observed for integers in the hippocampus (*p* = 0.86) and parahippocampal cortex (*p* = 0.57), though both regions showed a small effect for fractions (respectively *p* = 0.003 and *p* = 0.04).

When analyzing separately each of the three main math-responsive regions, only the IPS showed a positive and statistically significant correlation between fraction and integer magnitudes (integers: r = 0.28, *p* = 0.0001; fractions: r = 0.68, *p* = 0.0001; across the two domains r = 0.24, *p* = 0.0001). Although within-domain correlations were significant in MFG and pITG (MFG: integers: r = 0.37, *p* = 0.0001; fractions: r = 0.42, *p* = 0.0001; pITG: integers: r = 0.32, *p* = 0.0001; fractions: r = 0.58, *p* = 0.0001), cross-domain correlations were not (MFG: r = −0.20; temporal: r = −0.08).

Furthermore, within the parietal lobe, a continuous cortical map from larger to smaller numbers was observed, overlapping with areas PF and IP2 of the HCP atlas. As shown in figure 7B and C, the spatial patterns were broadly similar for fractions and integers, with smaller numbers represented in the most anterior part of the IPS, extending into the inferior parietal lobule, and increasingly larger numbers represented increasingly posteriorly in the horizontal part of the intraparietal sulcus. In these areas, consistent tuning was observed both within domain across the two independent runs (integers: r = 0.44, *p* = 0.0001; fractions: r = 0.91, *p* = 0.0001), as well as across the two domains (r = 0.69, *p* = 0.0001). Furthermore, a topographic map was confirmed by a significant correlation of the voxel tuning with the Z (inferior-superior) and X (left-right) coordinates of the MNI atlas, both for integers (Z: left *r* = 0.21, right *r*= 0.41; X: left *r* = 0.61, right *r* = −0.31) and for fractions (Z: left *r* = 0.83, right *r* = 0.70; X: left *r* = 0.56, right *r* = −0.58). In contrast, correlations with Y coordinate (anterior-posterior direction) were weaker (Y: left *r* = −0.34; right *r* = 0.08) and fractions (Y: left *r* = −0.36; right *r* = −0.30).

Finally, as a further proof of cross-domain consistency, when selecting voxels preferring numbers 0, 1, 2, or 3 in the integer map, we found that these voxels exhibited magnitude preferences consistent with fractional quantities. For example, voxels most responsive to 0 in the integer map also showed a higher activation for the fractions 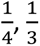, and 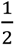 (Figure S2C).

The above results were obtained at the group level. At the single-subject level, although there was considerable variability across participants, the same general topographic pattern in the aIPS/IPL was observed in 14 participants for integers and 15 participants for fractions (at R^2^ > 0.3). Maps for fractions and integers from four representative participants are shown in supplementary Figure S3.

## Discussion

The ability to understand non-symbolic number magnitudes and ratios is present in babies (Izard et al., 2009; McCrink & Wynn, 2007), is shared with other species (Dehaene, Dehaene-Lambertz, et al., 1998; Vallentin & Nieder, 2008), and has been consistently mapped in the brain (Harvey et al., 2013; Lasne et al., 2019; Piazza et al., 2004). Yet, the symbolic systems that extend these foundational abilities, allowing us to represent novel concepts such as fractions and the symbol zero, reflect a uniquely human cognitive achievement. Despite their central role in mathematical reasoning, the neural mechanisms underlying symbolic numbers and their extension to such higher-level mathematical concepts remain much less understood. To investigate this, we used high-field fMRI during a number comparison task mixing integers and fractions to explore how symbolic numbers are represented at both behavioral and neural levels. Behaviorally, we observed the classic distance effect: participants were slower and less accurate when comparing numbers that were closer in magnitude, regardless of their domain (integers or fractions). Neurally, regions within the math network encoded number magnitude in a graded fashion: numbers closer in magnitude evoked more similar fMRI activation patterns, while those farther apart produced more dissimilar patterns. This pattern was also observed in the hippocampus and parahippocampal cortex. The neural representational geometry revealed an organization of neural activity along two distinct dimensions: one representing numerical domain, and another ordering numbers according to their magnitude along a cortical number line. Individual voxels showed reliable preferences for specific integers and fractions, and these preferences formed an organized mesoscopic “numerotopic” map in the bilateral aIPS/IPL, with similar voxel-tuning preferences for fractions and integers, indicating that these numerical domains are integrated into a single semantic map. Thus, the aIPS/IPL appears to play a key role in representing the magnitude associated with a symbolic number, whether it is a fraction or an integer.

Fractions differ from integers in several fundamental ways that could impact how they are mentally represented: 1) unlike integers, they are not organized according to successor relations; 2) an infinite number of fractions exist between any two numbers; and 3) their magnitude is not transparently related to the size of their component numbers (for instance, although 5 is greater than 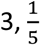 is smaller than 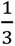). All these properties make fractions counterintuitive and difficult to learn, interfering with the formation of a mental number line for fractions in children (Cauté & Dehaene, 2025; Ischebeck et al., 2009; Schneider & Siegler, 2010). Nevertheless, our behavioral and brain-imaging findings indicated that, by adulthood, fractions and integers are well integrated into a single semantic space, at least within the small range we tested (0-5). These findings suggest that, to perform the task, participants transformed fractions and integers into a joint representation of their magnitude, thus mapping them onto a shared mental number line. Thus, by adulthood, in our educated participants, both evolutionary older concepts of integers and more recent and culture-dependent concepts of fractions have become integrated into a single representational map.

This conclusion fits with several prior behavioral and brain-imaging studies, which indicate that, by 5^th^ grade, the quantity meaning of fractions begins to be grasped and organized onto a mental number line (Binzak & Hubbard, 2020; Kalra et al., 2020; Park et al., 2025; Toomarian & Hubbard, 2018), though this development may be more protracted in French children (Cauté & Dehaene, 2025). Further converging evidences comes from Jacob and Nieder (2009), who used an adaptation paradigm to show that participants habituated to a given fraction (e.g., 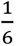), when subsequently presented with a fraction in numerical or word format (e.g., 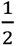 or “one-half”), showed a recovery of fMRI activity in the bilateral anterior IPS that varied as a function of the numerical distance between the deviant and the adaptor fraction. Our conclusion contrasts, however, with a single fMRI study suggesting that the neural code for fractions differs from that for integers (DeWolf et al., 2016). Closer examination, however, indicates that this is only a superficial discrepancy. Like ours, this study found a greater activation of the entire math-responsive network to fractions than to integers (DeWolf et al., 2016, Figure 4), and unlike us, it did not evaluate the fine-grained representational similarities of magnitudes across those domains. Furthermore, while our task required explicit comparisons of the magnitudes of fractions and integers, DeWolf et al. did not require direct comparisons between them, but presented integers, decimals and fractions in different fMRI runs, thus allowing participants to rely on domain-specific strategies (e.g., comparing the distance between denominator and numerator) rather than translating fractions into their numerical magnitude (Toomarian & Hubbard, 2018).

Our findings align with previous findings that the bilateral IPS is a key site where numerical magnitude is represented independently of notation (Bugden et al., 2019; Castaldi et al., 2020; Eger et al., 2003, 2009; Ischebeck et al., 2009; Jacob & Nieder, 2009; Mock et al., 2018; Piazza et al., 2007; Pinel et al., 2001; but see also Bulthé et al., 2014, 2015; Damarla & Just, 2013). Like others (e.g. Harvey & Dumoulin, 2017; Karami et al., 2025), we also found that magnitude information is present within the entire math-responsive network, also including bilateral hippocampus and parahippocampal cortices. This raises an important question: if the aIPS/IPL suffices to translate a number into its associated internal quantity, why do other core areas of the math network, as well as the medial temporal lobe, also appear to represent information according to its magnitude? A possible explanation is that once number symbols are translated into a numerical magnitude, this information may be broadcast to other regions forming a “global workspace” to support decision making and other working-memory dependent mathematical manipulations (Chateau-Laurent & VanRullen, 2025; Dehaene, Kerszberg, et al., 1998). Since fMRI is unable to decisively determine the direction of information flow in this network, an important future study would be to investigate the temporal dynamics of symbol comprehension and access to magnitude information using magnetoencephalography (MEG) or intracranial recordings (Pinheiro-Chagas et al., 2018, 2019, 2024).

Interestingly, we found that the hippocampus and parahippocampal cortex also represent information about integers and fractions in a graded manner, both within and between domains. Recent studies suggest that magnitude information may be encoded in the medial temporal lobe (Kutter et al., 2023; Menon, 2016), together with many other conceptual dimensions of words beyond the math domain (Franch et al., 2025). The hippocampus has been proposed to play a critical but time-limited role in the early phases of knowledge acquisition, by laying out the explicit structure of the learned domain (Behrens et al., 2018; Garvert et al., 2017), before this knowledge gets transferred to the cortex during skill development, as less efficient explicit procedures gradually giving way to more automatic and memory-based approaches (Baram et al., 2024). This may help explain why hippocampal engagement is often observed during arithmetic in children, but is less prominent in adults (Cho et al., 2012; Menon, 2016). The strong hippocampal involvement, in our study, may reflect the fact that fractions remain a challenging domain for many adults. An interesting avenue for future studies would be to investigate whether participants with different levels of mathematical education show systematic differences in how the hippocampus is recruited during numerical processing.

In all the cognitive maps that we inferred, from behavior (Figure 1), fMRI PCA (Figure 3B), fMRI RSA (Figures 4 and 5) or direct observation of cortical tuning to specific numbers (Figure 7), the neural representational geometry we observed aligned with previous research in showing that the concept zero occupies a position at the beginning of the mental number line (Barnett & Fleming, 2024) and that fractions are encoded according to their genuine magnitude (e.g., 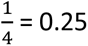), rather than to their numerator or denominator (e.g., 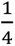 was far from 4) (Toomarian & Hubbard, 2018). The nature of our task, interleaving fractions and integers, likely prompted participants to compare the two domains by relying on a single number line (Toomarian & Hubbard, 2018) rather than applying distinct normalization strategies to each domain (Nelli et al., 2023; Sheahan et al., 2021;

Summerfield et al., 2020). Note that, in both behavior and math-responsive areas, the representational geometry inferred from MDS of the RSA matrix did not form a perfect straight line, but instead traced a one-dimensional curve embedded in two-dimensional space and where, furthermore, numbers 3-5 were close together and not always perfectly ordered (Figures 1 and 5; but see Figure 4). One possible explanation for the latter finding is that, due to the limited set of targets that we could study, the fractions ranged only from 0.25 to 2.5, thus covering only the smaller integers 0-3. This may have made all decisions on numbers 3-5 easier, thus causing an apparent bend in the values above three. Future experiments should ensure that fractions cover the full magnitude range of integers, making it more likely to observe a well-ordered number line. As for the curved pattern, a similar curved structure has previously been observed during a variety of ordering tasks in the parietal cortex and prefrontal cortex in humans (Barnett & Fleming, 2024; Karami et al., 2025; Karami, Alireza et al., 2026; Luyckx et al., 2019; Nelli et al., 2023) as well as in macaques (Okazawa et al., 2021) and is therefore not specific to our study. Curvature may allow the representational space to encode more than one dimension at once, with projection on one axis decoding stimulus order while projection on another orthogonal axis decodes stimulus difficulty and/or uncertainty (in our case, distance from the extremes 0 and 5) (Nelli et al., 2023). Several authors also warn against the possibility that this curvature or “horseshoe” pattern could be an artifact of MDS, which attempts to preserve the global distances between all pairs of items and, when it cannot respect those constraints in few dimensions, bends the linear continuum (de Leeuw, 2011; Diaconis et al., 2008; Gurnee* et al., s. d.; Proix et al., 2023). This possibility is all the more real that we found a one-dimensional organization when using the simpler method of principal component analysis (Figure 3B). Identifying the exact representational geometry of the cortical number line is an important problem for further research, which may ultimately require neuronal population recordings.

To the best of our knowledge, this is the first study to investigate the geometry of neural representations in a task involving both integers and fractions, as well as to identify a magnitude-based voxel-tuning preference map for both domains in the aIPS/IPL. By fitting a Gaussian function (Harvey & Dumoulin, 2017), we found that voxels indeed have consistent and cross-domain preferences for specific numbers (Figure 7A), as found at the single-neuron level in monkeys and humans (Kutter et al., 2018, 2023, 2024; Nieder & Dehaene, 2009). Furthermore, in one region (the anterior IPS extending towards the inferior parietal lobule), these preferences were organized according to a mesoscopic cortical map where voxels preferring small numbers are in the front and those preferring larger numbers occupy a more posterior and dorsal position (Figure 7B). This map overlaps with the horizontal segment of the intraparietal sulcus, an area implicated in numerical magnitude manipulation and whose damage leads to impairments in calculation abilities (Dehaene et al., 2003). While such topographic maps have been ubiquitously identified in response to the numerosity of non-symbolic concrete sets (Harvey et al., 2013; Harvey & Dumoulin, 2017; Paul et al., 2022; Tsouli et al., 2021), only a single study, focusing only on integers 1-7, uncovered tuning for symbolic numbers (Cai et al., 2023) which, surprisingly, was significant only in temporo-occipito cortex (at a site slightly posterior to the present pITG activation). In the present study, several factors may have facilitated the detection of symbolic tuning. One is the comparison task, which encouraged access to a magnitude representation. To perform the task, participants had to focus on the magnitude of the numbers, which is only one of the dimensions of number concepts (Dehaene et al., 2025). Thus, this magnitude information was likely amplified or expanded, as previously observed for other concepts (Çukur et al., 2013). In contrast, other studies used passive viewing paradigms or tasks that did not require explicit access to the numerical quantity, likely resulting in weaker magnitude representations, especially when probed with symbolic numerals. Furthermore, those studies, inspired by classic fMRI retinotopy paradigms, used a cyclic paradigm where numbers very progressively increase from 1 to 7 and back (Cai et al., 2023; Harvey et al., 2013; Harvey & Dumoulin, 2017). Thus, consecutive numbers were highly predictable and often repeated, which may have led to fMRI adaptation (Piazza et al., 2004) and participant boredom. In the present study, all transitions between numbers appeared equally often.

Another important factor may be task difficulty: comparisons involving only small integers are relatively easy, and the resulting neural activation may be weak. In contrast, by involving a mixture of fractions and integers, and by imposing a continuous load (since each number had to be kept in working memory in order to be compared to the next one), our sequential comparison task forced participants to continuously think of the magnitude of each number while maintaining sustained attention throughout the task. Indeed, Budgen, Woldorff, and Brannon (2019) showed that when participants engaged in a challenging task that required explicit magnitude processing, such as addition with two digits, the bilateral anterior IPS was similarly engaged by symbolic and non-symbolic formats, an effect that was not observed during a color-matching task (Bugden et al., 2019). The relationship between numerical formats seems to be less evident when participants perform easy tasks that do not require explicit magnitude computation, such as deciding if two numbers are equal (Lyons et al., 2015). Finally, the use of a 7T fMRI (1.2 mm isotropic, compared to 1.75 mm isotropic for Cai et al. (Cai et al., 2023)) likely allowed us to detect finer-grained patterns of brain activity.

While integers are natural to humans, fractions and the symbol zero are cultural tools developed out of scientific and mathematical inventiveness. Our findings reveal that with education, once they become easy and intuitive, both fraction and integer concepts end up being represented in terms of magnitude on the same internal cognitive and neural map. Furthermore, they provide particular clear support for the existence of cognitive maps (Gardenfors, 2000; Shepard & Chipman, 1970), i.e. geometric neural representations in which concepts with similar meaning are represented by similar vectors. High-field fMRI appears capable of monitoring, not only the large-scale cortical gradients that separate different categories of concepts (Deniz et al., 2019; Franch et al., 2025; Huth et al., 2016), but even the mesoscale representations that distinguish individual concepts within a given domain. With longer acquisition times, future work could extend the present findings to a broader set of math concepts (Debray & Dehaene, 2025) and clarify how cortical topography changes with education.

## Acknowledgements

We would like to thank Minye Zhan for her valuable assistance with the fMRI protocol, Manon Pietrantoni for sharing her PCA script, and all NeuroSpin support teams for their help in participant recruitment, stimulation, testing, data storage and analysis.

## Lead Contact

Further information and requests for resources should be directed to, and will be fullfilled by, the lead contact, Daniela Valério.

## Supplementary figures

**Figure S1.**
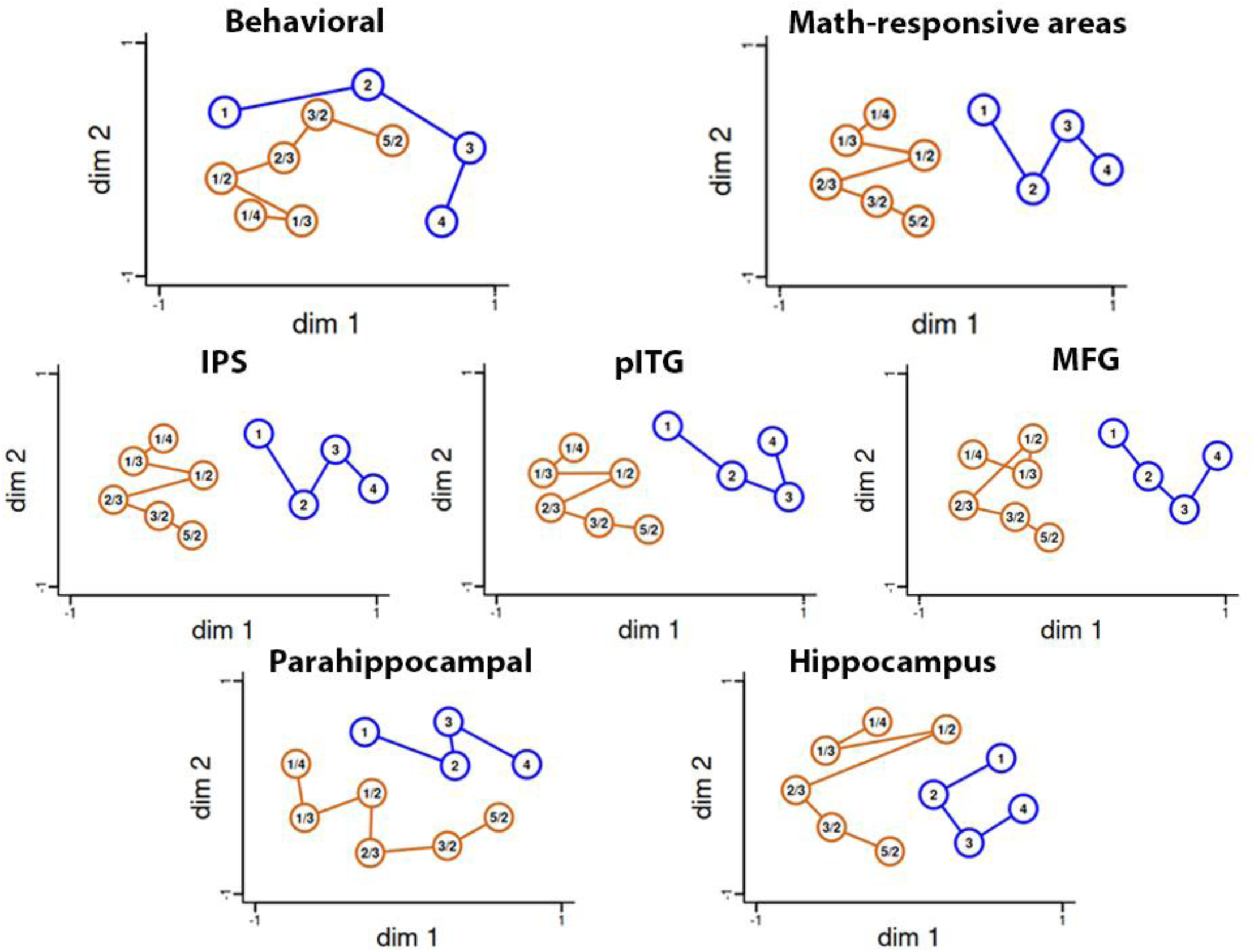
Two-dimensional MDS representation obtained without using the extreme numbers (0 and 5). MDS solutions were obtained using either the behavioral efficiency score matrix, math-responsive areas, as well as IPS, pITG, MFG, parahippocampal, and hippocampus.

**Figure S2.**
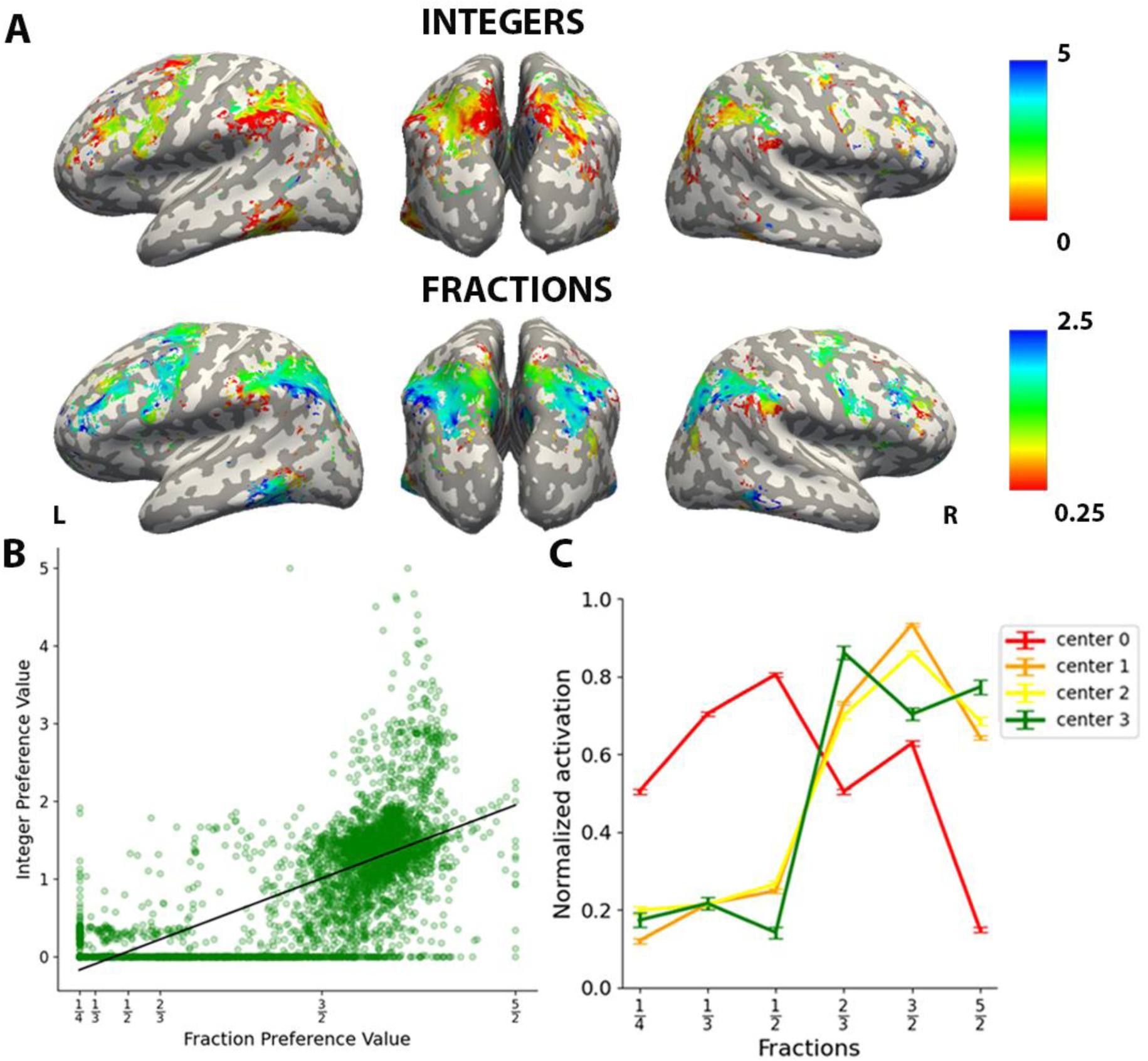
Further evidence for voxel tuning to specific numbers. A) Whole-brain view of the voxel tuning preferences for integers and fractions, with all math-responsive regions in the sentence localizer. Overall, the correlation between the two maps is very low (*r* = 0.01), but it is high and significant in the aIPS/IPL (*r* = 0.69) (panel **B**). Furthermore, in the aIPS/IPL, voxels preferring 0, 1, 2, and 3, selected based on their preferences in the integer map, show systematic changes in their normalized activation in response to fractions (panel **C**).

**Figure S3.**
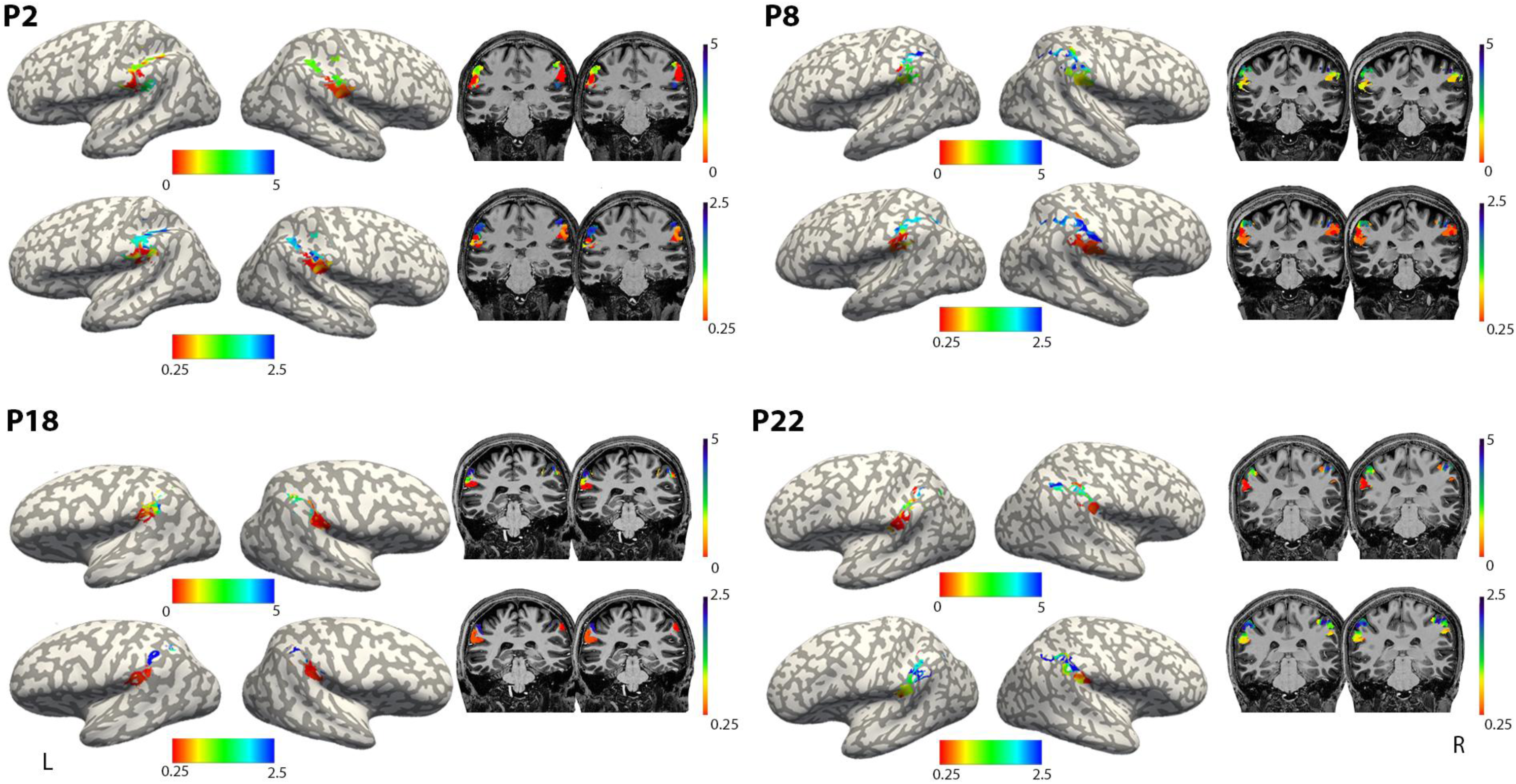
Single-participant maps of numerical preferences for integers (top) and fractions (bottom) in four participants.

**Table S1:** Effect of fraction-to-integer distance on neural dissimilarity matrices within the regions of interest used in the RSA (Figures 4 and 5). For each region, a multiple regression of RSA matrices was performed as described in Figure 6. The table reports the mean beta coefficient for the crucial fraction-to-integer distance predictor (averaged across subjects, with standard error of the mean in brackets), the peak MNI coordinates of the region, and the Z-score (representing a group-level one-sample t-test against zero).

| Region of interest | Average beta | MNI peak coordinates |  |  | Z score |
| --- | --- | --- | --- | --- | --- |
|  |  | x | y | z |  |
| Math areas | M = 0.33; SEM = 0.06 | 42 | -64 | -2 | 3.54 |
| IPS | M = 0.32; SEM = 0.06 | 61 | -42 | 47 | 3.57 |
| pITG | M = 0.29; SEM = 0.06 | 42 | -64 | -2 | 3.88 |
| MFG | M = 0.34; SEM = 0.06 | 28 | 40 | 23 | 3.49 |
| Hippocampus | M = 0.38; SEM = 0.06 | -24 | -37 | -8 | 3.32 |
| Parahippocampal | M = 0.40; SEM = 0.05 | -26 | -36 | -16 | 3.31 |

\*\* FDR, $q < 0.001$

